# Segmenting functional tissue units across human organs using community-driven development of generalizable machine learning algorithms

**DOI:** 10.1101/2023.01.05.522764

**Authors:** Yashvardhan Jain, Leah L. Godwin, Sripad Joshi, Shriya Mandarapu, Trang Le, Cecilia Lindskog, Emma Lundberg, Katy Börner

**Affiliations:** Department of Intelligent Systems Engineering, Luddy School of Informatics, Computing, and Engineering, Indiana University, Bloomington, IN 47408, USA; Department of Immunology, Genetics and Pathology, Division of Cancer Precision Medicine, Uppsala University, Uppsala, Sweden; Science for Life Laboratory, School of Engineering Sciences in Chemistry, Biotechnology and Health, KTH - Royal Institute of Technology, Stockholm, Sweden; Department of Bioengineering, Stanford University, Stanford, CA 94305, USA; Department of Pathology, Stanford University, Stanford, CA 94305, USA; Chan Zuckerberg Biohub, San Francisco, CA 94305, USA

## Abstract

The development of a reference atlas of the healthy human body requires automated image segmentation of major anatomical structures across multiple organs based on spatial bioimages generated from various sources with differences in sample preparation. We present the setup and results of the “Hacking the Human Body” machine learning algorithm development competition hosted by the Human Biomolecular Atlas (HuBMAP) and the Human Protein Atlas (HPA) teams on the Kaggle platform. We showcase how 1,175 teams from 78 countries engaged in community-driven, open-science code development that resulted in machine learning models which successfully segment anatomical structures across five organs using histology images from two consortia and that will be productized in the HuBMAP data portal to process large datasets at scale in support of Human Reference Atlas construction. We discuss the benchmark data created for the competition, major challenges faced by the participants, and the winning models and strategies.

## Introduction

The creation of a human reference atlas (HRA) requires harmonization and analysis of massive amounts of imaging and other data to capture the organization and function of major anatomical structures and cell types^1–3^. A key task is the segmentation of major anatomical structures—from the whole body to the single-cell level. Functional tissue units (FTUs) are used as a “stepping stone” from the organ to the single cell level. FTUs are defined as the smallest tissue organization that performs a unique physiologic function and is replicated multiple times in a whole organ. The spatial organization of FTUs matters and strongly impacts function. FTUs that are diseased have different cell type populations and possibly different sizes and shapes, or are altered in the number of FTUs within an organ. Several organ atlas efforts within the HuBMAP effort are now focusing on cell types, cell states, and biomarkers in specific FTUs. Being able to segment FTUs is an important part of identifying cell types and their gene/protein expression patterns within an FTU.

To segment anatomical structures in histological tissue sections efficiently, human intelligence must be efficiently combined with machine intelligence to overcome several challenges: segmenting histological images manually is labor-intensive, there are challenges with inter-observer variability, and there might be subtle differences and details that cannot be recognized or may be missed by the human eye. In support of efficient and high-quality tissue segmentation, human-in-the-loop approaches have been implemented^4,5^. Here, human expertise is used to identify and prepare relevant image data; design, optimize, train, and run effective machine learning (ML) algorithms; and interpret results. Once high-quality ML training datasets are compiled, generation and federation pipelines are set up, ML algorithms can be trained and optimized to segment image data at scale. As new datasets are segmented and these ML segmentations are validated and/or improved by human experts, ML algorithm performance can be further improved using this additional training data. Over the last decade, much work has been done on segmenting histological images; most of this work focuses on single cell segmentation^5^ or target structures in a single organ^4,6–8^, including functional tissue units (FTUs). FTUs are defined as the smallest level of tissue organization that performs a unique physiologic function^9^. To the best of our knowledge, there exist no ML algorithms that can segment FTUs across multiple organs in datasets from different laboratories.

In 2021, the Human BioMolecular Atlas Program (HuBMAP) conducted a Kaggle competition^10,11^ that focussed exclusively on segmentation of renal glomeruli in PAS stained histological images of kidney tissue, engaging 1,200 teams from 60 countries. The winning model from this competition was validated and productized in the HuBMAP data portal; it is now being run on all kidney tissue data at scale. In parallel, the Human Protein Atlas (HPA)^2^ conducted two Kaggle competitions^12–15^ that focussed on classification of subcellular patterns in cultivated cells in microscope confocal images, engaging nearly 3,000 teams across the two competitions. In addition to the confocal images of cultivated cells, the HPA has also generated >10 million immunohistochemically stained images from all major tissues of the human body^16^. HuBMAP and HPA partnered to address two major challenges when constructing a Human Reference Atlas (HRA): (1) standardization of data coming from various sources (different sample preparation and staining protocol, different equipment readout, etc.) and (2) robust and generalized segmentation of various tissue types. The two teams hosted a joint competition on Google’s ML community platform, Kaggle^17^, inviting competitors to develop machine learning algorithms that correctly segment FTUs of different shapes and sizes across five organs. This paper details the competition design (**Fig. 1**) and highlights the major challenges of the competition and the strategies used by the winning teams. We also present an analysis of competition dynamics and code performance improvements by 1,175 teams making 39,568 submissions over the 3- month period. Analogous to the previous HuBMAP competition, solutions from the winning teams will be productized in the HuBMAP data portal and ultimately process large amounts of relevant data to extract biologically relevant knowledge that can be used to construct the human reference atlas.

**Figure 1.**
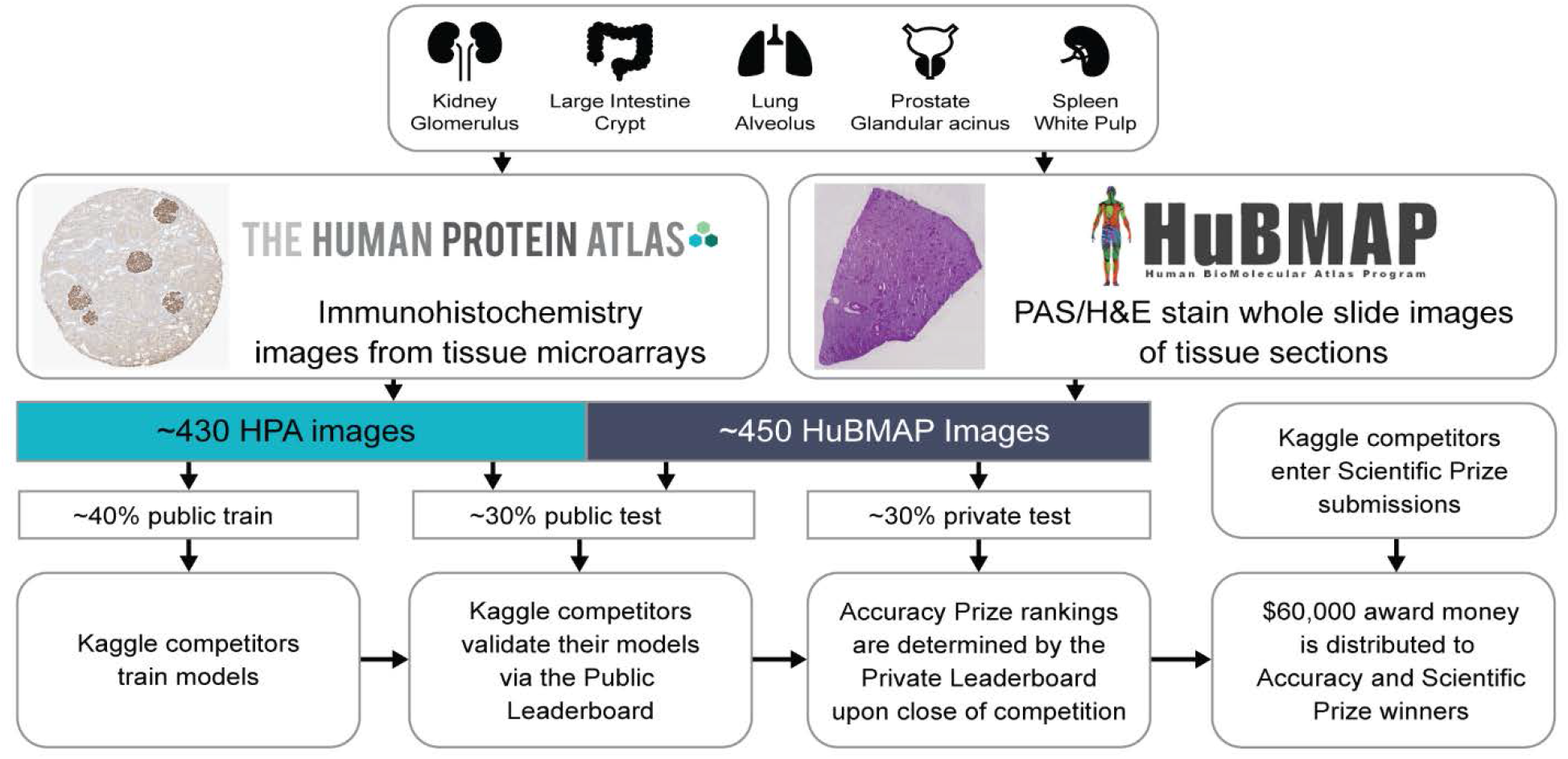
Overview of competition setup. Tissue data for five organs (top row) collected within HPA and HuBMAP using different tissue harvesting and processing protocols are collected and divided into a public training, public test, and private test dataset. Kaggle teams use the public training data to train their models; they then submit the models to the Kaggle Submission Portal to receive performance scores computed using the public test data. At the end competition, when all teams submitted their best algorithm solutions, all solutions were run against the private test set to determine Performance Prize winners.

## Results

### Competition Design and Performance Criteria

The aim of the “Hacking the Human Body”^18^ competition was to develop machine learning algorithms for segmentation of functional tissue units in five human organs using histology images sourced from two different consortia, namely HuBMAP and HPA (**Fig. 2**). The competition was designed to build algorithms that are generalizable across multiple organs and robust across dataset differences such as image resolution differences, color differences, artifacts, staining differences etc. HPA’s primary interest in this competition is that models that can segment FTUs in tissue sections can pave the way for more quantitative analysis of the data generated for the Tissue Atlas section of the HPA, e.g., to understand differences in protein expression patterns within FTUs as donor sex, ethnicity, or age change, or comparison of expression patterns of different proteins in the same donors. Human Reference Atlas construction in HuBMAP and other consortia use FTUs to characterize local cell neighborhoods with well-defined physiologic functions; they are interested in capturing differences in FTU numbers, sizes, and shapes for different donor demographics in health and disease. Being able to segment FTUs in tissue sections in histology images is an important step for characterizing their morphology, cell types and gene/protein expression patterns.

**Figure 2.**
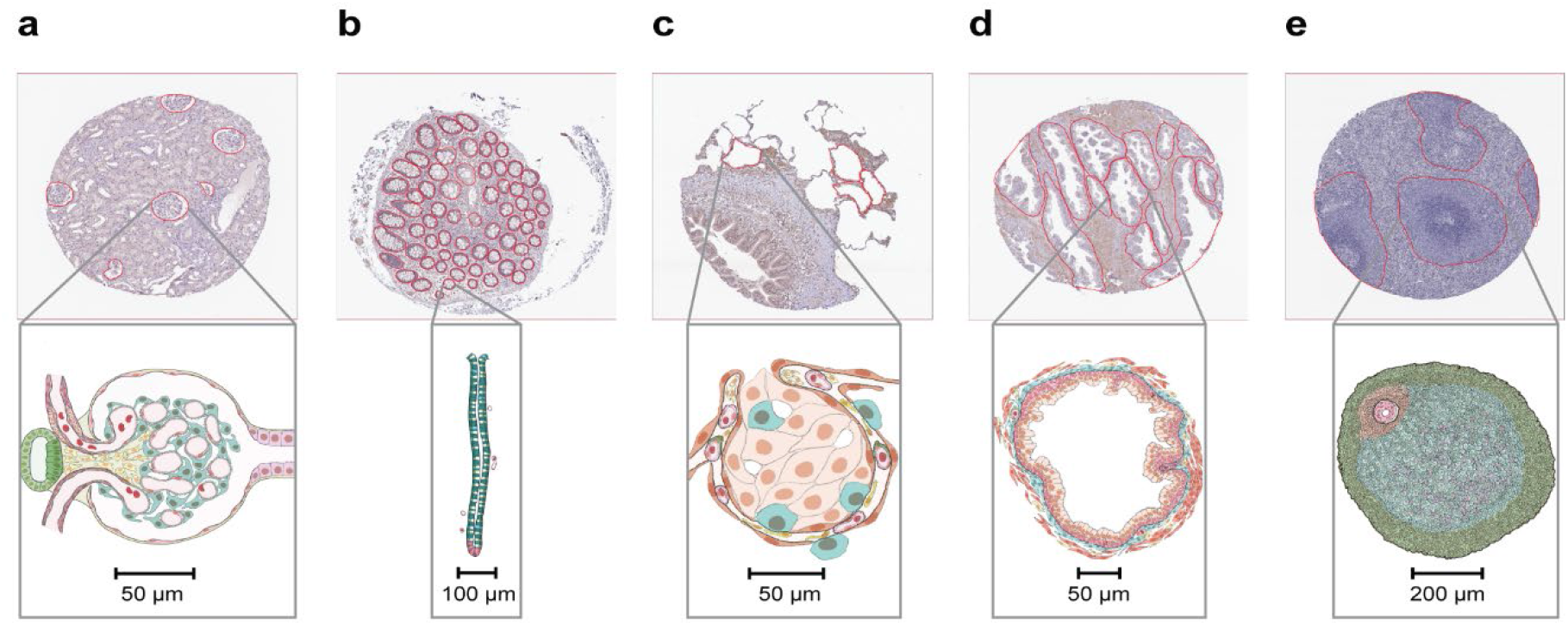
Exemplary tissue microarray cores with FTU segmentations outlined in red (top) and illustrations for all five FTUs (bottom). **a.** Glomerulus in the kidney. **b.** Crypt in the large intestine (top: perpendicular cross-section, bottom: lengthwise crosssection). **c.** Alveolus in the lung. **d.** Glandular acinus in the prostate. **e.** White pulp in the spleen.

The HPA and HuBMAP datasets cover five FTUs in five organs, namely renal glomeruli in kidney (renal corpuscle, UBERON:0001229), crypts in large intestine (crypt of Lieberkuhn of large intestine, UBERON:0001984), alveoli in lung (pulmonary alveolus, UBERON:0002299), white pulp in spleen (white pulp, UBERON:0001959), and glandular acini in prostate (prostate glandular acinus, UBERON:0004179). A dataset of 880 images was compiled, containing 432 images from HPA and 448 images from HuBMAP. This dataset was split into a training dataset of 351 images, and a private and public test dataset of 529 images (see **Table 1** for detailed breakdown). The HuBMAP dataset was preprocessed into a set of smaller tiled images (see Methods) to make HPA and HuBMAP datasets more comparable and to ensure teams could fully focus on developing machine learning algorithms rather than handling large format whole slide images (WSIs); providing image tiles also made the competition more equitable as computing requirements such as high RAM and high GPU access were not needed to develop code. Participants were allowed to use any external, publicly available data. All code submitted via the Kaggle submission portal was run on the public and private test set, leading to team rankings on the public and private leaderboards (see Methods). The algorithm performance is measured using the mean Dice coefficient on the test sets. The top-3 teams on the private leaderboard at the end of the competition win the performance prizes. Additionally, teams submitted entries to win the scientific prizes and the diversity prize (see Methods).

**Table 1.** Metadata for HPA data, both public and private, and HuBMAP data. All donors in the private HPA dataset are present in the public HPA dataset. All donors in the HuBMAP dataset are different from donors in the HPA dataset.

|  | # Images | # Male/Female<br>Unique Donors | Donor Age<br>Range | # FTUs |
| --- | --- | --- | --- | --- |
| **Public HPA Data** |  |  |  |  |
| kidney | 99 | 5/3 | 28-73 | 337 |
| large intestine | 58 | 3/4 | 47-84 | 3,107 |
| lung | 48 | 4/4 | 21-78 | 188 |
| prostate | 93 | 8/0 | 37-76 | 1,097 |
| spleen | 53 | 4/4 | 21-82 | 167 |
| **Public HPA Totals** | 351 | 24/15 | 21-84 | 4,896 |
| **Private HPA Data** |  |  |  |  |
| kidney | 19 | 4/3 | 28-70 | 69 |
| large intestine | 18 | 3/4 | 47-84 | 892 |
| lung | 14 | 3/4 | 43-78 | 66 |
| prostate | 18 | 7/0 | 37-76 | 212 |
| spleen | 12 | 3/3 | 21-74 | 38 |
| **Private HPA Totals** | 81 | 20/14 | 21-84 | 1,277 |
| **HPA Totals** | 432 | 29/17 | 21-84 | 6,173 |
| **HuBMAP Data** |  |  |  |  |
| kidney | 79 | 8/7 | 20-77 | 538 |
| large intestine | 43 | 2/2 | 22-48 | 1,966 |
| lung | 115 | 16/7 | 19-73 | 2,630 |
| prostate | 98 | 10/0 | 18-57 | 1,202 |
| spleen | 113 | 9/2 | 0-47 | 392 |
| **HuBMAP Totals** | 448 | 45/18 | 0-77 | 6,728 |
| **Totals** | 880 | 74/35 | 0-84 | 12,901 |

**Table 2.** Results for run time and accuracy from pilot run using baseline model.

| **Algorithm** | **Training Time (seconds)** | **Inference Time (seconds)** | **Mean Dice (Private HPA)** | **Mean Dice (HuBMAP)** | **Mean Dice (Private HPA + HuBMAP)** |
| --- | --- | --- | --- | --- | --- |
| Tom | 18,339 | 1,424 | 0.76 | 0.53 | 0.57 |

The major challenge in this competition was to build ML code solutions that are trained on one type of staining (from HPA) and can generalize to cover other types of staining (from HuBMAP) during inference. Consequently, teams developed strategies to deal with differences in terms of resolution, color, tissue thickness, etc. Additionally, teams had to optimize code for multiple organs, as lower performance on any organ would negatively impact the overall score. Other challenges included small training set size, uneven train/test split, and class imbalance, which motivated teams to build optimal solutions to extract maximum signal from the training data.

### Performance and Winning Strategies

The winning team for the performance prize reached a mean Dice score of 0.835 on the private leaderboard, followed closely by the second (0.833) and third (0.832) place winners. The score drops by 0.005 for the fourth place solution, reaching a mean Dice score of 0.827. The top-3 teams made a combined total of 619 submissions throughout the 3-month competition period. In general, the teams found kidney and large intestine to be the easiest classes, followed by spleen, prostate, and lung. Lung was the most challenging class in the competition (see **Supplementary Table 3**), primarily due to the variations in alveoli segmentations as they contained both collapsed and uncollapsed alveoli, as well as the variations in cuts (elongated vs. circular).

The main strategies that helped the teams to increase performance scores were data augmentation (geometric, color, distortions, scales) which involves artificially increasing the amount of data by adding many minor alterations to the original data, stain normalization (Vahadane method^19,20^), using external data for training, and pseudo labeling which involves adding increasingly confident label predictions from semisupervised training loop. All three winning teams used some version of all these strategies. Interestingly, the fourth place solution only used heavy stain normalization (reducing the importance of color in a model and forcing it to look for other cues in the images) and no external data or pseudo labeling, and was able to reach a mean Dice score of 0.827. Additionally, vision transformers proved to be extremely efficient compared to traditional convolutional networks due to their ability to capture long-range dependencies. However, such models are more sensitive to hyperparameter tuning and data changes. The teams found SegFormer^21^ models to be the best performing vision transformers. Since the SegFormer license is not completely open-source, teams also explored other vision transformer models and found Co-scale conv-attentional image Transformer (CoaT)^22^ models to be an effective replacement which performed equally well, while Swin^23^ transformers performed poorly. Finally, the second place solution showed that using bio-relevant auxiliary tasks such as organ classification and pixel size prediction also helps boost performance.

The first and third place winning teams also performed ablation studies (see **Supplementary Tables 1 and 2**) to assess the impact of different strategies on performance. The three most effective strategies were building ensembles of multiple models with at least one vision transformer model, using external data and pseudo labeling, and heavy data augmentation and/or stain normalization strategies. Team 3 used pixel size adaptation and histogram matching to heavily boost performance. Team 2 found that heavy encoders and networks with larger input resolutions worked better. Team 1 showed that while ensembles provide the best performance, the SegFormer mit-b4 model^21,24^ provides the best score (0.828) as a single model. This is an important result as ensembles are extremely resource intensive and can be impractical for some production settings. A single model combined with carefully selected image preprocessing strategies can be a good choice in production environments. Detailed code implementations and documentation of the three winning solutions can be found on GitHub (see Code Availability).

### Participation Analysis using Meta Kaggle

The competition ran from June 22, 2022 to September 22, 2022 and involved 1,517 individual competitors from 78 countries that collaborated in 1,175 teams. For 286 competitors, it was their first time participating in a Kaggle competition and 36 of them made the top-100 list on the private leaderboard. In total, the teams made 39,568 submissions and created 922 public code notebooks. Additionally, the participants created 224 public discussion forum posts and made 943 discussion comments. These metrics help understand the truly collaborative and globally inclusive nature of Kaggle competitions where teams interact extensively to share code, data, and knowledge.

Kaggle ranks all its users in five performance tiers based on their achievements and engagement on the platform, using their user Progression System^25^. In this competition, we had 22 Grand Masters, 103 Masters, 372 Experts, 559 Contributors, and 450 Novices participating (performance tier data is missing in Meta Kaggle for 11 users). The top-2 winning teams included experienced software engineers with a passion for machine learning and computer vision. The team winning third place consisted of computer scientists, machine learning researchers and analysts. Many participants came from biomedical backgrounds as well and shared their domain expertise generously via the discussion forums.

**Fig. 3** graphs the dynamics of the three-month competition. **Fig. 3a** shows the number of teams and messages and the progression of top leaderboard scores over the competition period. Note the sudden increase in the number of messages after the team merger deadline and winner announcement. The scores reached nearly 0.80 midway through the competition after which improvements were made through fine tuning solutions using techniques such as pseudo labels, using ensembles of multiple models, etc. Importantly, the public and private leaderboard scores remained similar throughout the competition leading to minimal shake-up at competition end and indicating a good dataset split between public test and private test datasets. **Fig. 3b** plots the performance tiers of participating users. **Fig. 3c** plots the number of submissions versus the highest private score; many of the 1,175 teams have few submissions with rather low scores; some teams have many submissions with high scores. Performance winners submitted 264 times (1^st^ place), 100 times (2^nd^ place), and 255 times (3^rd^ place).

**Figure 3.**
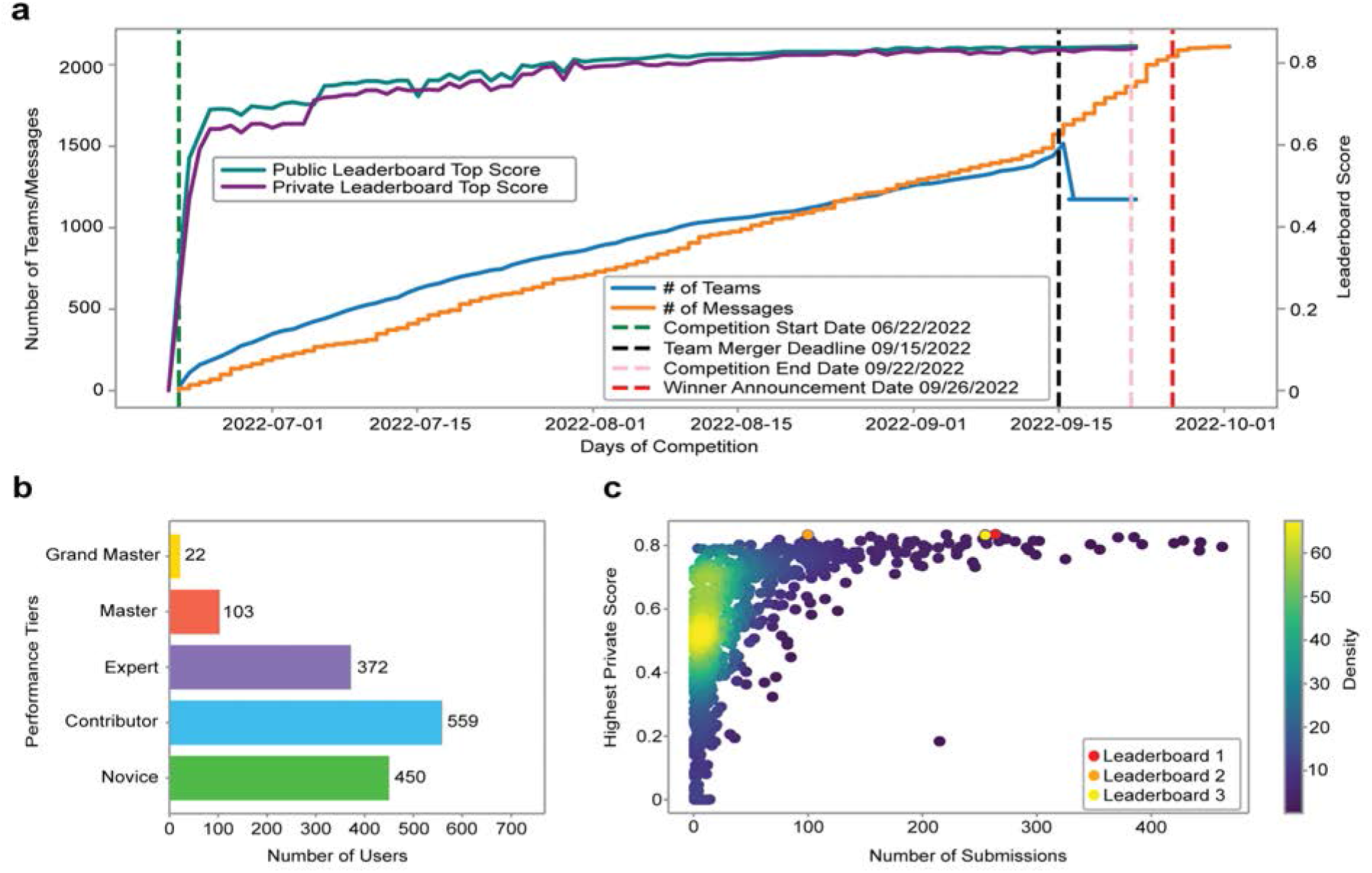
Competition dynamics over three months. **a.** Number of teams and messages and leaderboard high scores per day over competition period. **b.** Number of users by performance tier. **c.** Number of submissions vs. highest private leaderboard score for each of the 1,175 teams as a heatscatter.

### Scientific and Diversity Prizes

A total of six teams submitted their entries for the Scientific and Diversity prizes using a Google Form. The ten judges reviewed all submissions and graded them based on the rubric, ranking all submissions. Submission 5 and Submission 6 received the most points from all the judges for the two scientific prizes. Submission 5 focussed on showcasing differences between a convolution model and a vision transformer model, the latter achieving better performance as their bigger receptive field helps analyze images in a global context which is more suitable for medical image segmentation tasks. Additionally, it also showcased the importance of stain normalization in bridging the domain difference between the HPA and HuBMAP data. Submission 6 showcased the impact of noisy labels in the ground truth for training data and proposed a method to dynamically relabel missing annotations and minimize the gap between noisy and clean labels, thereby boosting performance by 4% on the private leaderboard.

All judges unanimously agreed Submission 1 should receive the Diversity and Presentation prize for building a team of diverse members and presenting their experiments and results in an extremely well documented and accessible manner.

## Discussion

Building the Human Reference Atlas is an extremely challenging task that requires close collaboration by experts from different scientific domains to solve key data integration, modeling, and visualization challenges across spatial and temporal scales. Kaggle’s open-source and community-driven nature makes it possible to bring in experts from industry, academia, government; to try out algorithms that were originally developed for different application domains; and to discuss solutions and results publicly empowering many to develop innovative solutions. All data and code is shared openly as a benchmark for use in future algorithm performance exercises and comparisons. Kaggle and other code competition platforms make it possible to share the burden of effective data preprocessing; run and compare thousands of ML algorithms in a very short period of time; and build on and advance these solutions collectively; something that is not possible at this speed and scale if research is performed in individual labs.

The “Hacking the Human Body’’ competition showcases the value of vision transformers in biological image processing, with all three winning teams building model ensembles consisting of some or all vision transformer models. This is in stark contrast to the previous HuBMAP competition^11^ (concluded in May 2021) where all winning teams used only convolutional models, evincing the quick rise of transformer models in the field. Sourcing ground truth labels for supervised learning tasks, especially in biomedical domains, is an extremely time-consuming and expensive challenge. The participants used diverse techniques to overcome this challenge, including using additional unlabeled data and creating pseudo-labels for training iteratively to improve performance using a semi-supervised approach. This, in addition to clever data augmentation and normalization techniques, turned out to be the key to building generalized solutions that can be deployed at scale.

While this competition provides several innovative and high-performing solutions, there exist several limitations of these models for real-world production use cases: 1) Since the models are trained on a small dataset, there is risk of model overfitting; 2) The vision transformer models, as teams realized via many iterations of experiments, are much more sensitive to hyperparameter tuning and data changes than the convolutional models; 3) Model ensembles are computationally expensive and might be impractical or inefficient for many production environments.

Going forward, we plan to address the above-mentioned limitations by training and validating the models on more data, and compressing the large ensembles into single models. The code from the winning models will be productized and deployed in the HuBMAP data portal to process large amounts of tissue data and extract biological knowledge in support of Human Reference Atlas construction and usage.

## Methods

### Competition and Prizes

The “HuBMAP + HPA - Hacking the Human Body” competition was conducted on Google’s ML and data science community platform called Kaggle, from June 22, 2022, to September 22, 2022. The private leaderboard for identifying the three performance prize winners was finalized on September 26, 2022. The Judges prize winners were announced 3 weeks later, after a thorough review and discussion by the judges panel. The winners of the performance prize were awarded with a cash prize (US$15,000 for first place; US$10,000 for second place; US$5,000 for third place). The winners of the Judges’ prize were also awarded with a cash prize (US$10,000 each for two scientific prizes; US$10,000 for one diversity prize).

#### Performance Prize

A fast and efficient performance evaluation metric was required to score hundreds of submissions per day and a total of 39,568 over three months. The teams submitted their inference code, after training their models, on the Submission portal. The submitted code was then run over the public test set to rank the teams on the public leaderboard. The teams typically use this score to validate their models. The teams can make an unlimited number of submissions before the competition deadline, limited to 5 submissions per day. On competition end, the teams can choose up to two solutions to submit as their final submissions, which are then scored on the private test set, which always remains inaccessible to the teams, to rank the teams on the private leaderboard. All scoring is done using the mean Dice score as the evaluation metric (see Evaluation Metrics under Methods) and the top-3 teams on the private leaderboard are selected as winners for the performance prize.

#### Judges’ Prize

Judges’ prizes were aimed to promote experimentation, diversity, and science communication. The scientific prizes aimed to motivate solutions that go beyond the Dice evaluation metric and are more experimental in nature, providing insight into the dataset and/or computational techniques. The diversity and presentation prize promoted inclusion and the effective communication of scientific results. The winners were determined by a panel of human judges using a predefined and publicly available evaluation rubric (see **Supplementary Information**) that was publicly available on the Kaggle competition website at competition start.

### Dataset Collection and Assembly

All tissue data used in this study is from donors examined and identified by pathologists as pathologically unremarkable tissue that can be used to derive the function of healthy cells. As the focus of this work is on the identification of FTUs, all images used in this competition feature at least one FTU.

#### HPA Data

The HPA data consists of immunohistochemistry images of 1 mm diameter tissue microarray cores and 4 μm thickness, stained with antibodies visualized with 3,3’- diaminobenzidine (DAB) and counterstained hematoxylin (H)^16,26^. We retrieved over 7TB of public data from the HPA which comprised 23,610 images of 1 mm diameter tissue microarray (TMA) cores for the large intestine, 27,906 for kidney, 28,098 for lung, 28,934 for prostate, and 27,474 for spleen. Given that the HRA aims to capture human adults, we removed all images associated with patients below the age of 18. We computed sex, age, tissue region percentage per image and selected 500 public images that maximize sex and age diversity per organ, have at least 1 FTU, and have a tissue region percentage above a threshold value (where threshold value is 5% for lung and 15% for kidney, spleen, large intestine, and prostate). The resulting dataset has 432 public HPA images distributed across the five organs (**Table 1**). We further retrieved about 44GB of private (unpublished) data from the HPA which comprised 253 images for large intestine, 295 images for kidney, 291 images for lung, 265 images for prostate, and 290 images for spleen. This dataset was processed in the same way as the public HPA data. A total of 81 images were selected for the final private dataset. All images, both public and private, are 3,000 px x 3,000 px (with some exceptions as roughly 19 images lie between 2,308 x 2,308 px and 3,070 x 3,070 px), and the diameter of each tissue area within an image is approximately 2,500 px x 2,500 px which corresponds to 1 mm. Hence, the pixel size of the images in this dataset is roughly 0.4 μm.

#### HuBMAP Data

Multiple teams within or affiliated with HuBMAP shared 257 periodic acid-Schiff (PAS)^27^ or hematoxylin and eosin (H&E)^28^ stain WSIs of healthy human tissue that has not been published. From these WSIs, 1mm x 1mm tiles were extracted to match the size of the HPA TMA core images. Minimum donor metadata for all WSIs used in this competition included organ name, sex, and age. The resulting dataset had 448 image tiles distributed across the five organs and sourced from five different teams. All donors across all organs were above the age of 18, an exception being spleen which included younger donors of ages 0 through 18. The pixel size of images across different organs was 0.5 μm for kidney, 0.229 μm for large intestine, 0.756 μm for lung, 0.494 μm for spleen, and 6.263 μm for prostate. The tissue slice thickness of all images in HuBMAP data was between 4-10 μm: 10 μm for kidney, 8 μm for large intestine, 4 μm for spleen, 5 μm for lung, and 5 μm for prostate.

#### Dataset Sampling

Some images feature space without human tissue. We calculated the tissue region percentage for each image using Otsu’s^29^ thresholding. The specific threshold values for each organ were selected manually by analyzing the number of images available against different threshold values (**Supplementary Figure 1**). The values were selected such that images with very low actual tissue area are discarded, yet leaving a sufficient number of images to work with.

We then constructed a dataset with similar numbers of donor samples across age groups and sex for all organs (**Supplementary Figure 2a**)--insofar possible given available HuBMAP and HPA data. Note that systematic sampling of healthy human organ tissue is non-trivial; while human donors do not mind giving up adipose tissue, getting tissue from other organs is often only possible if an organ transplant cannot be executed or a patient dies and the tissue is released for single-cell research. Consequently, the number of donors above the age of 50 is higher than those below 50, especially for the HPA data (**Supplementary Figure 2b**).

#### Data Format

For consistency, all images are exported as TIF files and all segmentations are provided as run-length encoded (RLE) masks for efficient storage and submissions (along with original JSON files) to the teams. Note that the RLE versions of the segmentation masks are cleaner than the JSON masks, although differences are minor. The JSON versions are more raw and the annotations might have issues, like overlaps, that do not exist in the RLE copies but can also allow the teams to distinguish between multiple adjacent FTUs which would all end up in the same mask with RLEs.

### Acquiring Ground Truth Labels and Final Dataset

For four organs (except kidney), 1-3 trained pathologists and/or anatomists (with experience in segmentation and histology) per organ provided initial segmentations done manually. For the kidney, the winning model from the previous HuBMAP Kaggle competition was used to generate initial FTU segmentations for all HPA and HuBMAP kidney data which were then manually reviewed and corrected by a professional anatomist.

All segmentations were verified and corrected through a final expert review process conducted by the lead pathologist for each organ. All images that were considered unsuitable were rejected. Partial FTUs were accepted, provided a human expert can segment it with confidence. All annotators, during the initial segmentation process as well as during the final review process, were given access to the images via an internal web-based segmentation tool (originally developed by the HPA team and further modified by the HuBMAP team). Please note that while extreme care was taken to get the best possible ground truth segmentations from experts, the labels do contain some noise, due to human bias, and existing issues were openly discussed on the public discussion forums of the competition.

#### Final Dataset

The final dataset used in the competition contains 432 images from the HPA (including 351 public and 81 unpublished images with a total of 6,173 FTU annotations) and 448 unpublished images from HuBMAP (with a total of 6,728 FTU annotations). All data is divided into three distinct datasets: a public training dataset containing all public HPA data (351 images), a public test dataset containing all unpublished HPA images (81 images) and HuBMAP images (209 images), and a private test dataset containing only HuBMAP images (239 images) (**Fig. 1**). The training dataset is openly accessible to the teams, while the test datasets remain hidden.

### Baseline Segmentation Model

To ensure the task is neither too easy (i.e., nearly 100 percent accuracy is achieved within little effort) nor too hard or impossible to accomplish (i.e., a satisfying accuracy is impossible), initial runs using the winning algorithm from the first HuBMAP Kaggle competition, Tom, created a baseline model. The model was run on Indiana University’s Carbonate large-memory compute cluster, using the GPU partition which consists of 24 Apollo 6500 GPU-accelerated nodes where each node is equipped with two Intel 6248 2.5GHz 20-core CPUs. We used a single node with 300 GB of RAM and 2 Nvidia V100- PCIE-32GB GPUs.

The model required about 5 hours for training and nearly 20 minutes for the inference task. It achieved a mean Dice score of 0.76 and 0.53 on the private HPA data and HuBMAP data, respectively. The mean Dice value achieved across the total private test dataset (HPA and HuBMAP) is 0.57. The same model achieved a mean Dice value of about 0.95 for the task of segmenting renal glomeruli in kidney images in the first HuBMAP Kaggle competition. The results of this baseline model demonstrate the task is neither too easy nor too difficult, and there is a need for more generalizable algorithms.

### Evaluation Metrics

The metric used to rank the performance of the teams in the competition is mean Dice coefficient (also referred to as the mean Dice score). The Dice score compares the pixel-wise agreement between a predicted segmentation (PS) and its corresponding ground truth (GT) segmentation for an image: 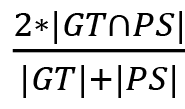.

The leaderboard score used is the mean of the Dice coefficients for each image in the test set. It should be noted that calculation of Dice coefficient does not take into account separation between individual instances. Hence, in case multiple predicted FTUs overlap/merge, the Dice coefficient for that prediction may still be high while the FTU count may be incorrect (and might require further processing, either programmatic or manual, to separate the individual instances of FTUs).

After extensive discussion of options with the Kaggle data scientists and machine learning experts from the panel of judges, the mean Dice coefficient was selected for performance prize ranking. While other metrics such as the mean Average Precision (mAP) might have been better suited for the problem, the Kaggle team recommended going forward with the mean Dice score, taking into account the nature of the dataset and timeline for the competition. Dice is a well-tested metric used in many competitions on the Kaggle platform and other metrics require much more testing by the Kaggle team to ensure participants cannot find loopholes and exploit vulnerabilities in the metric during the competition. Hence, while Dice score may not be the ideal metric^30,31^ in a production setting, it is a good enough metric to evaluate and compare solutions from Kaggle competitions.

### Public and Private Leaderboards

Kaggle ranks teams on two leaderboards–public leaderboard and private leaderboard–each using a different subset of the test data, using the predetermined evaluation metric for the performance prize. The public leaderboard uses the public test data and the private leaderboard uses the private test data. The public leaderboard rankings and scores are visible to the teams and are used to validate their algorithms, providing feedback they can use to improve their algorithms. The private leaderboard rankings remain hidden to the teams until the end of the competition to ensure algorithms are not overfit to test data. The top-3 teams on the private leaderboard are considered as winners of the performance prizes.

### Participation Analysis

At the conclusion of the competition, participation metadata becomes publicly available on Meta Kaggle^32^–Kaggle’s public data on competitions, users, submission scores and kernels. Meta Kaggle tables were initiated in 2015 and are updated daily with information on completed competitions. We use this data to understand how the “Hacking the Human Body” competition unfolded over its 3-month period.

We use standard python packages for data science such as Pandas^33^, NumPy^34^, Matplotlib^35^, and Seaborn^36^ for running all analyses; creating all visualizations in Jupyter^37^ Notebooks. The analyses can be replicated for any competition on Meta Kaggle using the code we made available on GitHub (see Code Availability).

## Acknowledgments

We appreciate the generosity of the sponsors of this competition: Google HCLS and Genentech (a member of the Roche group). We thank Andrea de Souza (Eli Lilly and Company) for obtaining sponsors. We thank Amy Kemper, Sohier Dane, and Addison Howard (Google/Kaggle team) for expert support throughout the competition.

We thank the HPA personnel Mattias Forsberg and Kalle von Feilitzen (both from Royal Institute of Technology) for assistance in making the images accessible to the challenge and HuBMAP teams for providing data and segmentation expertise for five organs: Jeff Spraggins (VU), John Hickey (Stanford University), Gloria Pryhuber (URMC), Doug Strand (UTSouthwestern), Maigan Brusko (UFL), Sanjay Jain (Washington University School of Medicine in St. Louis), Jeanne Shen (Stanford University), Iain Miller (Stanford University), Benjamin Dulken (Stanford University), Gail Deutsch (University of Washington). Mike Gallant (Indiana University) and Jason Swedlow (University of Dundee) kindly provided assistance for transferring and compiling the HPA data. Bruce *W*. Herr II (Indiana University) helped provision the web-based segmentation tool. Naveksha Sood (Indiana University) helped run code for generating pre-segmentations on HPA and HuBMAP data.

We are grateful to the Kaggle Scientific and Diversity prize judges Zorina Galis (NIH), Carolina Wählby (Uppsala University), Artem Sokolov (HMS), Constantin Kappel (Leica Microsystems), Anna Kreshuk (EMBL), Blue Lake (UC San Diego), David Van Valen (CalTech), Jhimli Mitra (GE Research), Nathan Heath Patterson (VU), and Bobak Kechavarzi (Cleveland Clinic) for sharing their time and expertise.

This research has been funded in part by the NIH Common Fund through the Office of Strategic Coordination/Office of the NIH Director under award OT2OD026671 and OT2OD033756, NIH awards U54EY032442-01 and U54HG010426-01, National Institute of Diabetes and Digestive and Kidney Diseases (NIDDK) award U54DK120058, the NIDDK Kidney Precision Medicine Project grant U2CDK114886, and the Knut and Alice Wallenberg Foundation. The content is solely the responsibility of the authors and does not necessarily represent the official views of the National Institutes of Health.

## Data Availability

All data used in the competition, along with the trained models, is publicly available on GitHub https://github.com/cns-iu/ccf-research-kaggle-2022.

## Code Availability

All code used for data preprocessing and analysis, baseline model, and winning algorithms, are publicly available on GitHub https://github.com/cns-iu/ccf-research-kaggle-2022.

## Author Contributions

YJ aided the design and implementation of the competition; led data analysis, sampling and processing; refactored and maintained web-based segmentation tool; aided segmentation and review process by experts; oversaw the competition throughout its duration; acted as liaison to data providers, segmentation experts, HPA team and Kaggle Team; co-wrote the paper. LLG aided the design and implementation of the competition; led data and metadata collection from HuBMAP data providers; aided data segmentation and review process by experts; acted as liaison to data providers, segmentation experts, HPA team and Kaggle Team; implemented initial code for participation analysis and visualizations; co-wrote the paper. SJ contributed to data analysis, sampling, and processing; conducted baseline model training and analysis. SM ran participation analysis and rendered the resulting visualizations. TL aided implementation of the competition; helped using the web-based segmentation tool; performed data and metadata assembly from HPA. CL aided design and implementation of the competition; guided data and metadata assembly from HPA; led the pathology-based data generation of the original HPA dataset. EL aided design and implementation of the competition; led data and metadata assembly from HPA. KB led the design and implementation of the competition; co-wrote the paper.

## Supplementary Information

### Supplementary Figures

**Supplementary Figure 1.**
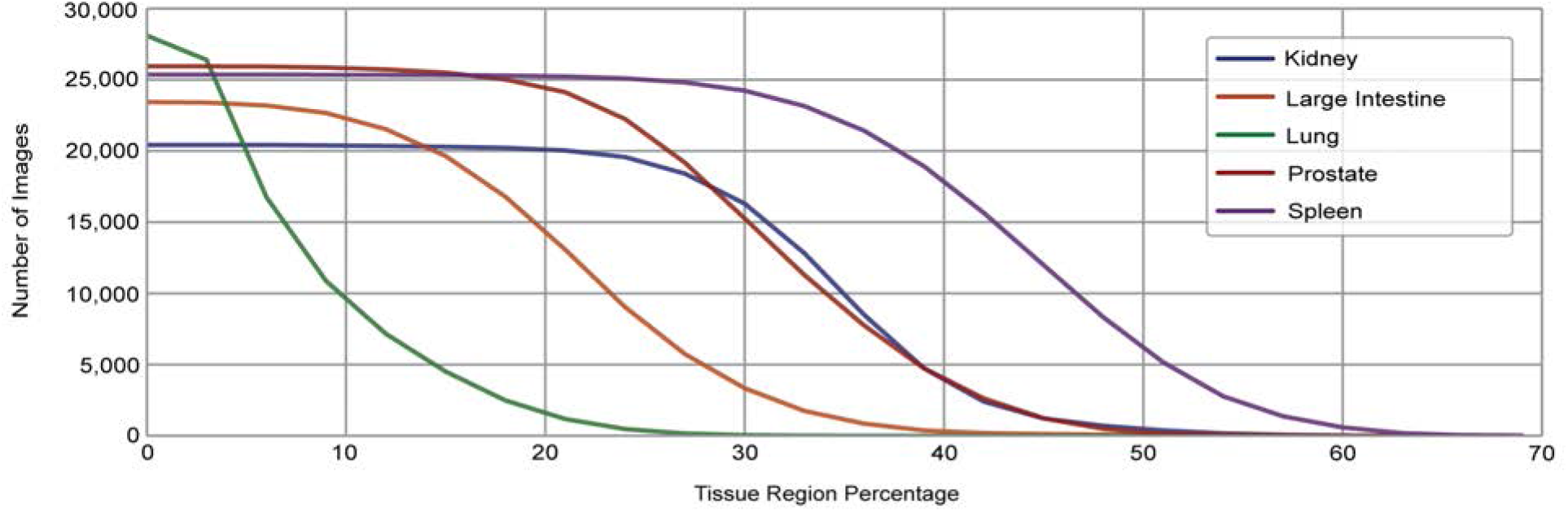
Tissue region percentage vs. number of images against different threshold values for public HPA data for all five organs. Final selected tissue region percentage thresholds: 5% for lung and 15% for other four organs.

**Supplementary Figure 2.**
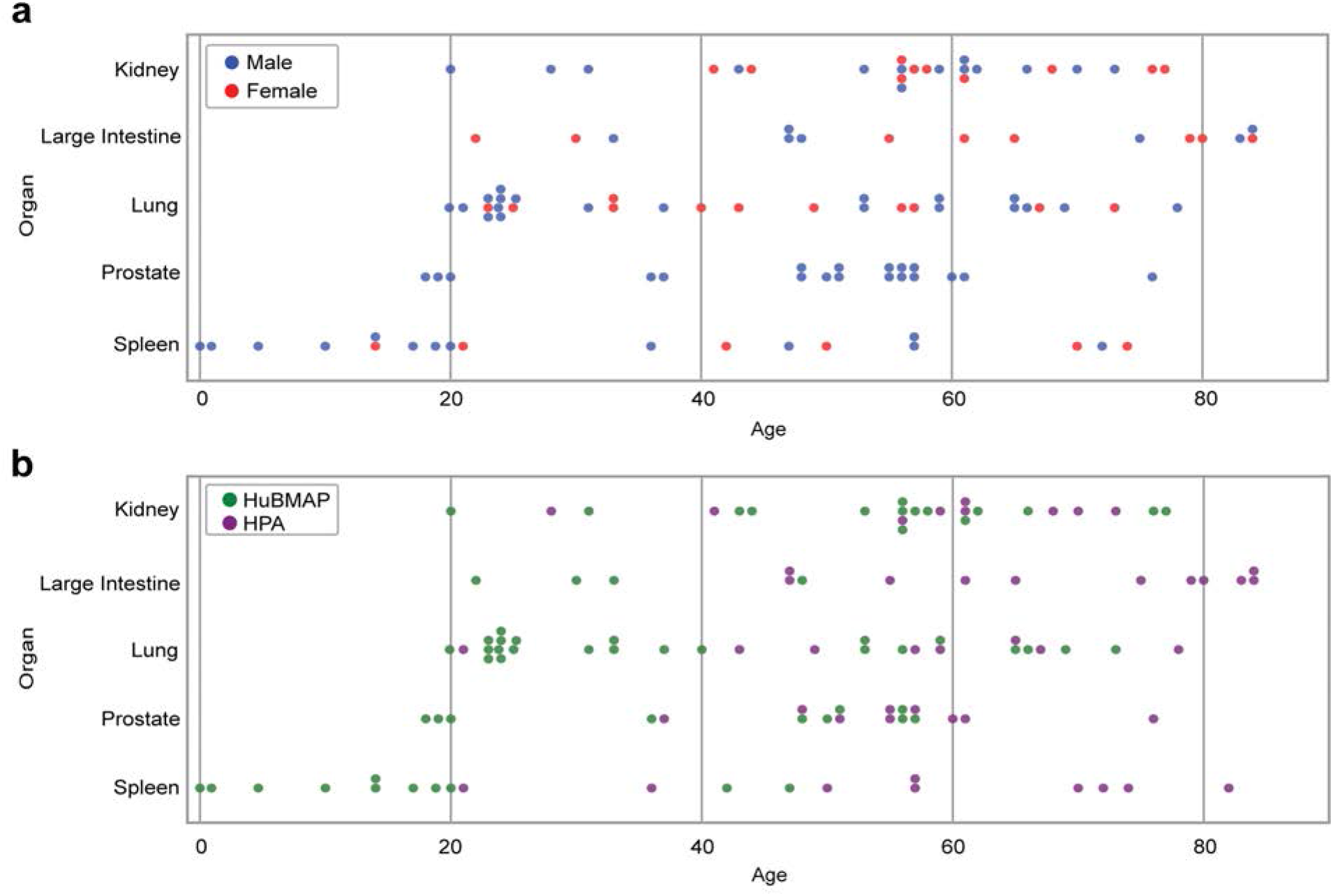
Tissue samples per organ and age for all 5 organs. **a.** Donor distribution color coded by male (blue) and female (red). **b.** Donor age and sex distribution color coded by HPA (purple) and HuBMAP (green).

### Supplementary Notes

#### Supplementary Notes 1. Judges’ Prizes Rubric

##### 1.1 Scientific Prize

Kaggle teams were asked to investigate correlations between predicted FTUs (e.g., area and shape) and donor demographics (e.g., sex and age). The evaluation rubric further emphasized validation of methods and implementations, documentation of performance and limitations, novelty of solutions, and presentation of insights useful for generation of reference FTUs for inclusion into a Human Reference Atlas. Scientific Prize winners were identified by a panel of human experts who selected two teams to receive equal Scientific Prize amounts ($10,000 each) based on the submission’s contribution to science and demonstration of innovative approaches. The complete evaluation rubric, as presented below, consisted of eight criteria which were used by the judges to evaluate the winners. Each criterion consisted of ten points for a total of 80 points.

1. Are the statistical and modeling methods used to identify FTUs appropriate for the task?
2. Are confidence scores and other metrics provided that help interpret the results achieved by the segmentation methods?
3. Is the presented characterization of FTUs useful for understanding individual differences, e.g., the impact of donor sex and age on the shape, size or spatial distribution of FTUs?
4. Is it possible to predict FTU area size distribution, given age and sex information across all organs?
5. Did the team validate their methods and algorithm implementations and provide information on algorithm performance and limitations?
6. Did the team document their method and code appropriately?
7. Did the team develop a creative or novel method to segment FTUs?
8. Did the team provide insights that would be useful for generating reference FTUs for inclusion into a HUman Reference Atlas?

##### 1.2 Diversity and Presentation Prize

The complete evaluation rubric, as presented below, consisted of three criteria which were used by the judges to evaluate the winner. Each criterion consisted of ten points for a total of 30 points.

1. Does the team embrace diversity and equity, welcoming team members of different ages, genders, ethnicities, and with multiple backgrounds and perspectives?
2. Did the authors effectively communicate the details of their method for segmenting FTUs, and the quality and limitations of their results? For example, did they use data visualizations to present algorithm setup, run, results and/or to provide insight into the comparative performance of different methods? Were these visualizations effective at communicating insights about their approach and results to experts and novice users?
3. Are the important results easily understood by the average person?

#### Supplementary Notes 2. Alveoli segmentations in lung tissue

Due to confusion regarding varied looking alveoli segmentations in lung tissue images, additional information was provided to the teams. The data included masks of both atelectatic (collapsed) and inflated alveoli (un-collapsed). The alveolar appearance on the image slides depends on how the tissue samples were prepared. For the inflated alveoli, which have a 3D ‘cup’ shaped structure, how the tissue is sectioned can cause variability as well. If the alveoli were sectioned in a horizontal manner, their shape will appear more like a complete circle. Whereas if the alveoli were sectioned vertically, they may appear more as a U-shape.

### Supplementary Table Legends

All tables can be accessed at Supplementary Tables.

**Supplementary Table 1.** Team 1 Ablation Study. This table lists the ablation study done by the winning team, detailing the strategies that helped improve the performance of their solution.

| Model | Private Dice | Public Dice | Public Hubmap Dice | Private Dice Gain |
| --- | --- | --- | --- | --- |
| mit-b4 + lung annotation | 0.79318 |  | 0.58097 | 0.07256 |
| mit-b4 + prostate_downscale | 0.68486 |  | 0.52633 | 0.05791 |
| mit-b4 + albu dataaug + pseudo label(prostate, intestine) | 0.72062 | 0.76051 | 0.54628 | 0.02496 |
| mit-b4 + better resize | 0.69566 |  | 0.52913 | 0.0108 |
| mit-b4 + stain transfer(torchstain) | 0.80838 | 0.80858 | 0.59243 | 0.00927 |
| mit-b4 aug + brightness | 0.82451 |  | 0.6046 | 0.00853 |
| mit-b4 + external lung | 0.82054 | 0.81345 | 0.60057 | 0.00622 |
| mit-b4 + external spleen | 0.81432 |  | 0.59722 | 0.00594 |
| mit-b4 + trainval(351 images) | 0.79799 | 0.80006 | 0.58248 | 0.00481 |
| 3 model ensemble | 0.83238 | 0.82364 |  | 0.00417 |
| mit-b4_optimized_threshold | 0.82786 |  | 0.60663 | 0.00335 |
| 4 model ensemble + image ration divisor 32 | 0.8338 | 0.82622 |  | 0.00142 |
| mit-b4 + pseudo label(prostate, intestine) patch | 0.79911 |  | 0.58466 | 0.00112 |
| 6 model ensemble | 0.83562 | 0.82716 |  | 0.00094 |
| 5 model ensemble | 0.83468 | 0.82679 |  | 0.00088 |
| mit-b4_0.8_1_1.2 | 0.82821 |  | 0.60718 | 0.00035 |
| mit-b4 | 0.62695 |  | 0.47984 | 0 |
| mit-b4 aug + HPA original (not stained) | 0.81598 | 0.81661 |  | -0.00456 |

\* Sorted in increasing order of private gain.
\*\* List is non-exhaustive and does not include all experiments by the team. Only listed is the subset of experiments team tracked and provided.

**Supplementary Table 2:**
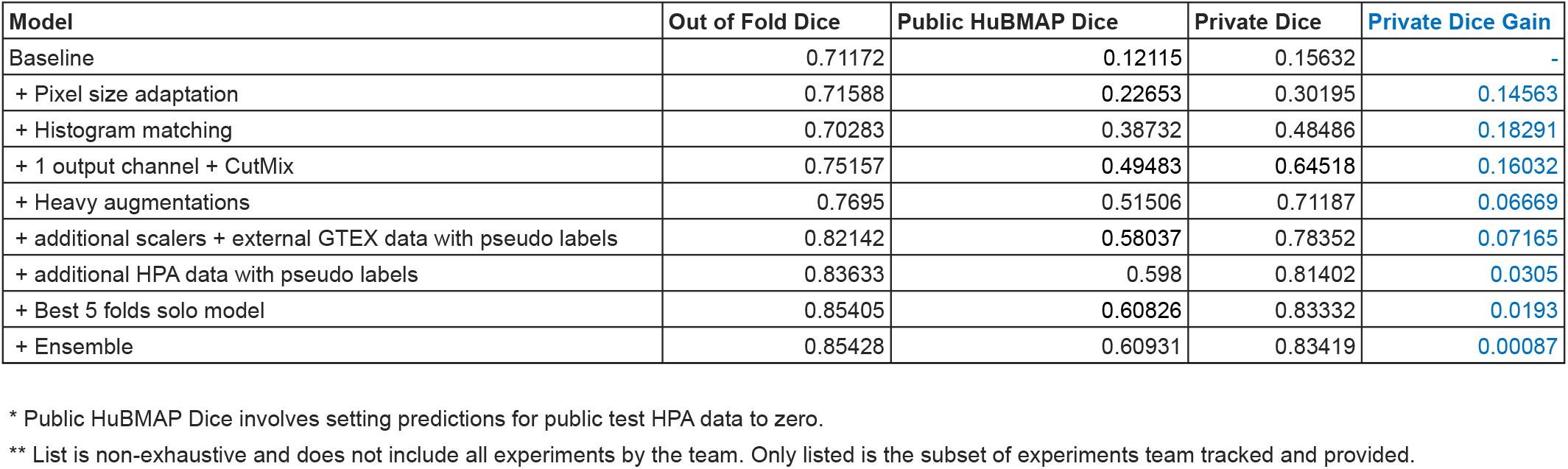
Team 3 Ablation Study. This table lists the ablation study done by the team that won the third performance prize, detailing the strategies that helped improve the performance of their solution.

**Supplementary Table 3:** Organ Dice Score Comparison for Teams Winning the Performance Prizes. This table provides the organ-wise breakdown of performance results for the three winning teams.

| **Team** | **Kidney** | **Large Intestine** | **Lung** | **Prostate** | **Spleen** | **Overall** |
| --- | --- | --- | --- | --- | --- | --- |
| First place winner | 0.96401 | 0.89676 | 0.72664 | 0.85004 | 0.83862 | 0.83562 |
| Second place winner | 0.9665 | 0.88931 | 0.72092 | 0.84851 | 0.84157 | 0.83393 |
| Third place winner | 0.9491 | 0.86232 | 0.73599 | 0.84806 | 0.84211 | 0.83266 |

\* All scores presented for private test set.

